# Mutational analyses reveal PLP-independent functions at PipY, the cyanobacterial paradigm for pyridoxal-phosphate binding proteins

**DOI:** 10.64898/2025.12.29.696868

**Authors:** Antonio Llop, Lorena Tremiño, Asunción Contreras

**Affiliations:** Departamento de Fisiología, Genética y Microbiología, Universidad de Alicante, 03690 San Vicente del Raspeig, Spain

**Keywords:** *Synechococcus*, PipX, B_6_ homeostasis, RNA binding, PLPHP

## Abstract

Pyridoxal-phosphate binding proteins (PLPBP) are involved in the homeostasis of B_6_ vitamers and amino/keto acids, share a high degree of sequence conservation and are represented in all three domains of life. Despite the obligate presence of the catalyst cofactor PLP, attempts to show enzymatic activity have been unsuccessful. Instead, evidence of RNA binding activity has been provided for several members of the family. Here we use PipY, one of the few PLBPB members studied so far, as a model system to address the phenotypic impact in the cyanobacterium *Synechococcus elongatus* of mutations K26A, P63L and R210Q, which respectively prevent PLP binding or are equivalent to those conferring B_6_-dependent epilepsy in humans with a recessive inheritance pattern. We found that while mutation K26A at the PLP-binding residue abrogated all phenotypes associated to PipY overexpression and toxicity, P63L and R210Q behaved as dominant gain-of-function mutations that inhibited bacterial growth. We provide *in vivo* evidence of PipY performing PLP-independent functions, in which mutant variant PipY^K26A^ but not PipY^P63L^ or PipY^R210Q^ would be defective. A model integrating our observations with previous data from other organims and PLPBP variants is discussed.

## Introduction

The Pyridoxal phosphate (PLP)-binding proteins (COG0325/PLPBP) are involved in vitamin B_6_ and amino acid homeostasis (Labella et al. 2017; Ito et al. 2019; Vu et al. 2020; Ito 2022; Tremiño et al. 2022). PLPBP/PLPHP (for PLP homeostasis proteins) family members are found in all kingdoms of life (Prunetti et al. 2016; Farkas and Fitzpatrick 2024) exemplified by the proteins YBL036C (yeast), YggS (Gram-negative bacteria), YlmE (Gram-positive bacteria), PipY (cyanobacteria) and PLPHP (previously called PROSC in humans or plants). They are all single domain proteins exhibiting the type-III fold of PLP-holoenzymes (Schneider et al. 2000; Eswaramoorthy et al. 2003; Tremiño et al. 2017; Tremiño et al. 2018; He et al. 2022) whose deficiency alters the B_6_ pool in the different systems studied (Darin et al. 2016; Prunetti et al. 2016; Ito et al. 2019; Johnstone et al. 2019; Ito et al. 2020; Vu et al. 2020). In humans, the severity of *PLPBP* mutations causing vitamin B_6_-dependent epilepsy appears to correlate with decreased ability of the PLPHP variants to bind the PLP cofactor (Johnstone et al. 2019) although only some of the pathogenic mutations found in patients complement the pyridoxine-sensitive phenotype of the *E. coli yggS* null mutant (Darin et al. 2016). In zebrafish the complete absence of PLPHP triggers epilepsy seizures and, importantly, early death, indicating the essentiality of the protein in development (Johnstone et al. 2019) and further suggesting an important regulatory complexity.

PipY from the cyanobacterium *Synechococcus elongatus* PCC7942 (hereafter *S. elongatus*) is one of the best-characterized PLPBP members and can be regarded as a paradigm for these proteins (Labella et al. 2017; Cantos et al. 2019; Tremiño et al. 2022; Llop, Labella, et al. 2023). In *S. elongatus*, *pipY* null mutants display PLP-related phenotypes including increased sensitivity to pyridoxine and to the antibiotics D-cycloserine and β-chloro-D-alanine (Labella et al. 2017). In addition, recombinantly produced PipY has been used to mimick the effect of selected pathogenic mutations on protein function (Tremiño et al. 2017). Fig. 1 illustrates the structural parallelism between cyanobacterial PipY and human PLPHP, highlighting the PipY residues Lys26, Pro63 and Arg210, whose mutations are analysed in this work.

**Figure 1.**
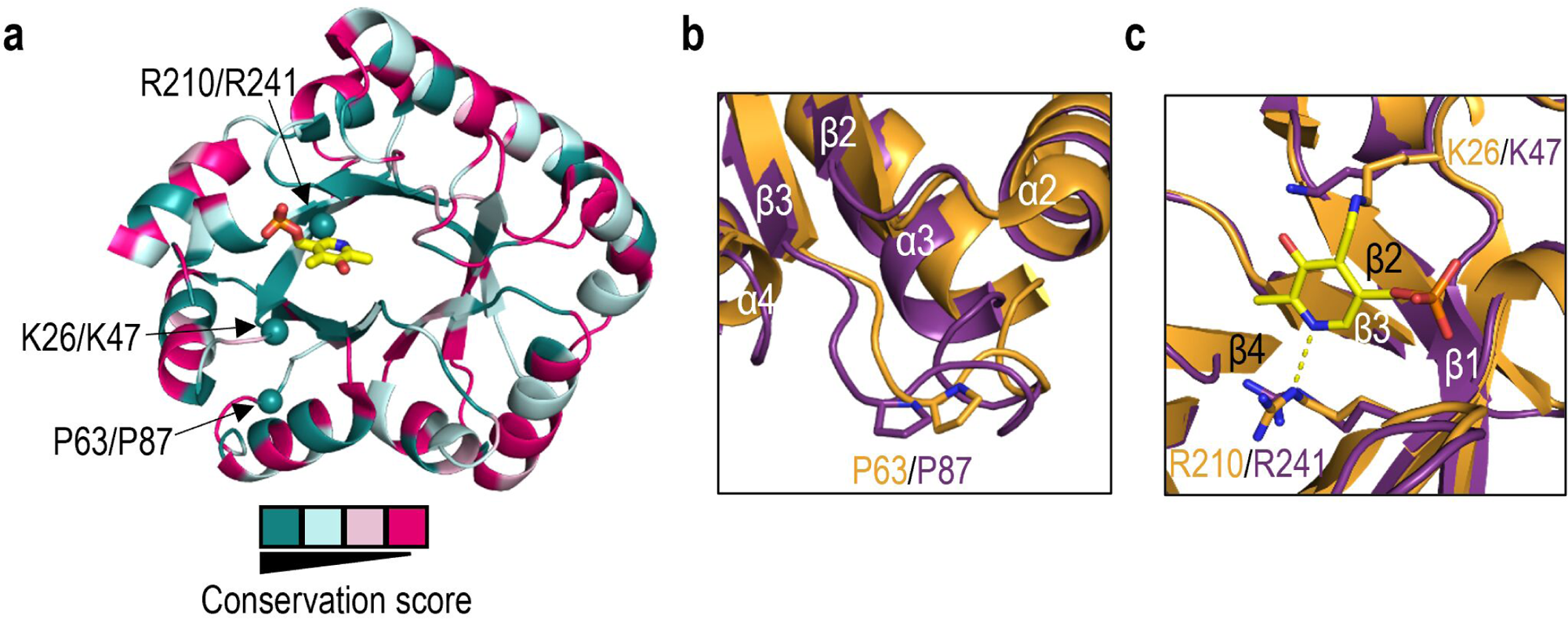
Conservation between cyanobacterial and human PLPBP orthologs and location of relevant residues mutated in this work. **a)** Structure of PipY from *S. elongatus* (PDB ID: 5NM8), colored according to the evolutionary conservation score of residues between PipY and its human ortholog (PLPHP). Color coding was defined based on the conservation score of an automatic ClustalW alignment of both protein sequences, as indicated in the legend. Relevant residues are mapped as spheres. PLP is shown in stick representation and labeled, with C, O, N, and P atoms colored yellow, red, blue, and orange, respectively. **b, c)** I-TASSER superimposition of the structural model of PLPHP over PipY colored purple and orange, respectively, with the residues of interest and secondary-structure elements indicated.

Despite the obligate presence of the catalyst cofactor PLP, no enzymatic activity has been associated to the PLPBP family and the current view is that they would have purely regulatory functions as components of signal transduction pathways (Tremiño et al. 2022; Graziani et al. 2024). In bacteria *PLPBP* genes cluster with cell division and cell wall biosynthesis (*dcw*, in Gram-positive bacteria and mycobacteria), PLP salvage, surface motility, secretion, amino acid metabolism, and translation genes (Prunetti et al. 2016; Tremiño et al. 2022). In cyanobacterial genomes the *sepF* gene (involved in cell division and restricted to gram-positive bacteria and cyanobacteria) is typically located downstream *pipY*. In *S. elongatus* and in most cyanobacteria *pipY* is part of a bicistronic *pipXpipY* operon with relatively short or non-existent intergenic distances suggestive of tight co-regulation and functional interactions between their products in this bacterial phylum (Labella et al. 2017; Cantos et al. 2019).

While deciphering the mechanisms of action and regulatory connections of PLPBP members remains very challenging, the inferred functional association between PipY with PipX provides a unique opportunity to investigate PLPBP functions in the context of a signalling pathway that is highly conserved in an important biological group of organisms. PipX, a hallmark of cyanobacteria, is a small protein involved in metabolic and environmental signalling that form complexes with other regulatory proteins, transcriptional regulators, or translation-related factors (Burillo et al. 2004; Espinosa et al. 2006; Espinosa et al. 2007; Llácer et al. 2010; Laichoubi et al. 2011; Espinosa et al. 2018; Labella et al. 2020; Jerez et al. 2021; Jerez et al. 2024; Salinas et al. 2024). PipX stabilizes the conformation of the transcriptional regulator NtcA and helps local recruitment of RNA polymerase (Forcada-Nadal et al. 2025) in response to nitrogen limitation (Espinosa et al. 2014; Giner-Lamia et al. 2017). PipX also interacts with the essential ribosome-assembly GTPase EngA (YphC/Der/YfgK) (Jerez et al. 2021; Llop, Bibak, et al. 2023). Both PipX and PipY regulate the levels of a common set of transcripts and afect the susceptibility to PLP-targeting antibiotics D-cycloserine and β-chloro-D-alanine, further suggesting their involvement in the same genetic pathway(s) (Labella et al. 2017).

Neither PipX or PipY are required for growth of *S. elongatus* under standard laboratory conditions (Espinosa et al. 2009; Labella et al. 2017; Cantos et al. 2019), although their overexpression or certain gain-of-function mutations at PipX prevent growth (Laichoubi et al. 2012; Labella et al. 2017; Jerez et al. 2021; Llop, Labella, et al. 2023). Overexpression of either PipX or PipY also induce a specific stress program termed chlorosis or bleaching (Jerez et al. 2021; Llop, Bibak, et al. 2023), a complex adaptative response by which they degrade their light-harvesting antenna, the phycobilisome, and reduce their chlorophyll content (Schwarz and Grossman 1998; Spät et al. 2018; Forchhammer and Schwarz 2019). PipY overexpression also confers specific phenotypic traits in *S. elongatus* including increase of cell length, decrease of cell viability and photosynthesis activity, accompanied by a dramatic and unprecedented accumulation of giant polyphosphate (hereafter polyP) granules (Labella et al. 2017; Llop, Labella, et al. 2023).

The aim of this work was to gain insights into yet unknown functions of the PLPBP family by taking advantage of the already existing structural and functional information on PipY. Here we address the phenotypic effects conferred in *S. elongatus* or *Escherichia coli* strains by PipY variants expressing point mutations K26A, P63L and R210Q that together prevent PLP binding or mimic pathogenic changes found in the orthologous human protein.

## Materials and Methods

### Plasmid construction

The plasmids and primers used in this study are listed in Table 1 and S1, respectively. Cloning procedures were carried out in *Escherichia coli* XL1-Blue. All constructs were verified by automated Sanger sequencing.

**Table 1.** Plasmids.

| Plasmid | Description, Relevant Characteristics | Source or Reference |
| --- | --- | --- |
| pUAGC127 | *pipY* replaced with *cat*, Ap<sup>R</sup> Cm<sup>R</sup> | (Labella et al. 2017) |
| pUAGC128 | *pipXpipY* replaced with *cat*, Ap<sup>R</sup> Cm<sup>R</sup> | (Labella et al. 2017) |
| pUAGC280 | (*cs3 lacI Ptrc*) into *NSI*, Ap<sup>R</sup> Sm<sup>R</sup> | (Moronta-Barrios et al. 2013) |
| pUAGC294 | (*cs3 lacI Ptrc:pipY*) into *NSI*, Ap<sup>R</sup> Sm<sup>R</sup> | (Labella et al. 2017) |
| pUAGC1129 | (*cs3 lacI Ptrc:pipY*<sup>628cga>caa</sup>) into *NSI*, Ap<sup>R</sup> Sm<sup>R</sup> | This work |
| pUAGC1130 | (*cs3 lacI Ptrc:pipY*<sup>199ccc>ctg</sup>) into *NSI*, Ap<sup>R</sup> Sm<sup>R</sup> | This work |
| pUAGC1133 | (*cs3 lacI Ptrc:pipY*<sup>76aag>gcg</sup>) into *NSI*, Ap<sup>R</sup> Sm<sup>R</sup> | This work |

Plasmids pUAGC1133, pUAGC1130, and pUAGC1129 carrying mutant alleles *pipY*^K26A^, *pipY*^P63L^, and *pipY*^R210Q^, respectively, were generated by QuickChange site-directed mutagenesis using pUAGC294 as template with primer pairs PipY-K26A-F/PipY-K26A-F, PipY-P63L-F/PipY-P63L-R and PipY-R210Q-F/PipY-R210Q-R, respectively.

### E. coli assays

Transformation of *Escherichia coli* XL1-Blue and MG1655 (Table 2) was performed by heat shock, essentially as described in (Froger and Hall 2007). 65 μL of competent cells were incubated with 50 ng of the plasmids. After the heat shock, 1 mL LB was added cells were incubated at 37°C for 1 hour with agitation. An equivalent volume to 100 or 900 μL of XL1-Blue and MG1655 cells, respectively, were spread on LB agar (1.5% w/v) plates. Ampicillin was added to solid media at a concentration of 75 μg/mL and, when indicated, the appropriate amount of IPTG. Pictures were taken after 24- or 48-hours incubation at 37°C.

**Table 2.**
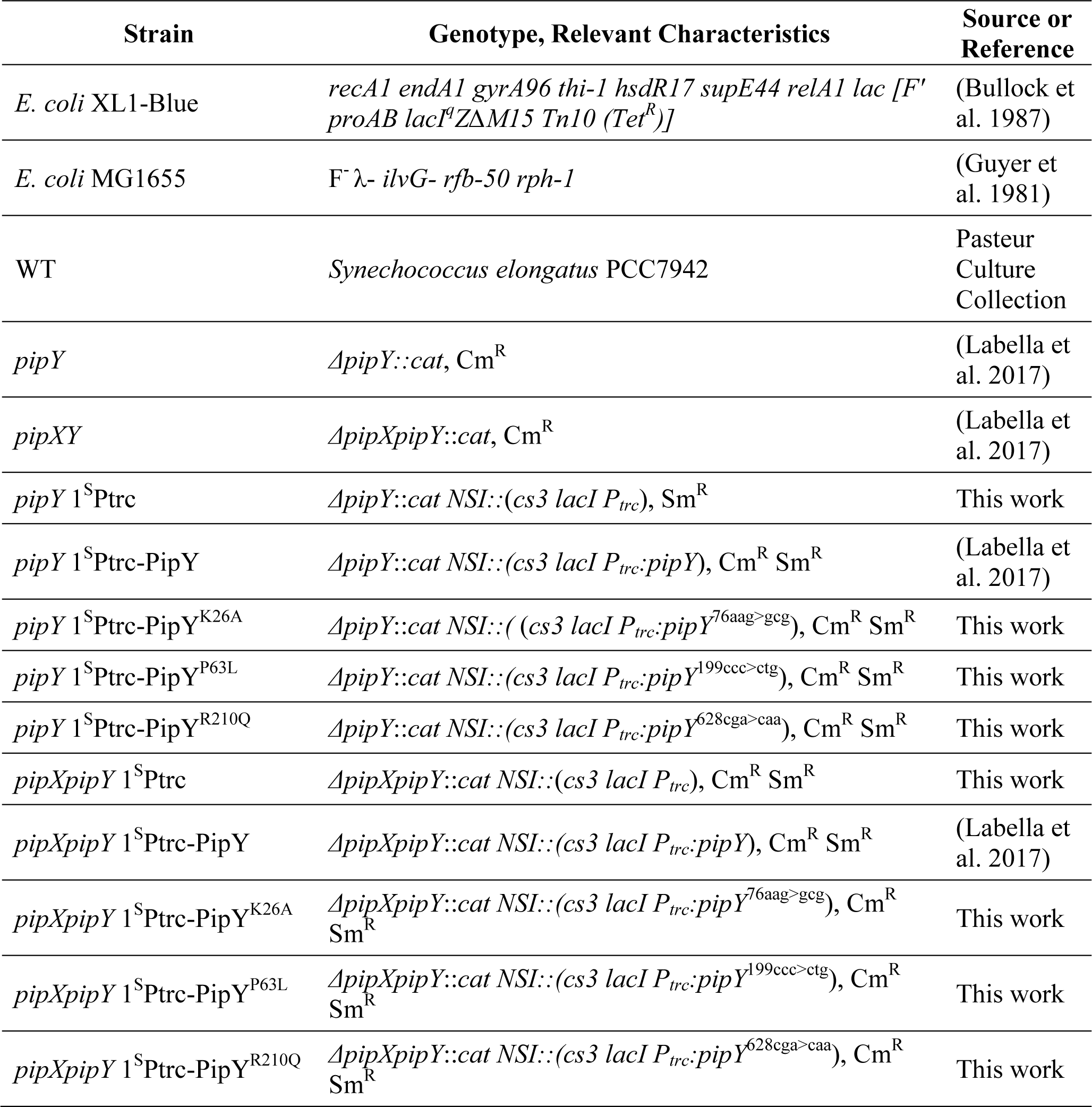
Strains.

### Cyanobacteria culture conditions and strain generation

*S. elongatus* cultures were routinely grown in blue–green algae BG11 medium (BG110 supplemented with 17.5 mM sodium nitrate (NaNO₃) and 10 mM HEPES/NaOH (pH 7.8; (Rippka et al. 1979)) at 30 °C under constant cool white fluorescent light, either in liquid cultures (150 rpm, 70 μmol photons m⁻²s⁻¹; mix of two clones) or on plates (50 μmol photons m⁻² s⁻¹; individual clones). Solid media contained 1.5% (w/v) agar and 0.5 mM sodium thiosulfate (Na₂S₂O₃). When appropriate, chloramphenicol (Cm, 3.5 μg/mL) or streptomycin (Sm, 15 μg/mL) were added.

For liquid growth, cultures of 30 mL were grown in BG11 using baffled flasks. Growth was monitored by measuring the optical density at 750 nm (OD_750nm_) in 1 mL samples using an Ultrospec 2100 Pro UV-Vis Spectrophotometer (Amersham Biosciences, Amersham, UK). All experiments were performed on mid-exponential phase cultures (OD_750nm_ = 0.4 – 0.8). To overexpress proteins the indicated concentration of isopropyl β-D-1-thiogalactopyranoside (IPTG) was added to media.

*S. elongatus* strains used in this study are listed in Table 2. Transformations were performed essentially as described in (Taton et al. 2020), and correct allele replacement verified by PCR. The primer pairs used were inter2060-1F/2059-Cm-1R and inter2060-1F/PipX-5R-129 for *pipY*, PipX-126-F/2060-Cm-1R for *pipXpipY*, and 7942NSIA-F/ NSI-1R for NSI.

### Determination of growth and pigment content

For growth measurements in flasks experiments, 25 mL of cultures were adjusted to an initial optical density at 750 nm (OD_750_) of 0.1 using a Ultrospec 2100 Pro UV-Vis Spectrophotometer (Amersham Biosciences, Amersham, UK). OD_750_ was then recorded at 0, 24 and 72 h after addition of 50 µM IPTG. For drop-plate assays in solid media, 5 μL of culture samples adjusted to an OD_750_ of 0.5 (and serial dilutions of 1/2, 1/10 and 1/100) were spotted on BG11 plates without or with 50 µM IPTG. Plates were incubated for 5 days under standard conditions.

### Cell staining and confocal microscopy parameters

To detect polyP granules in *S. elongatus,* 4′,6-diamidino-2-phenylindole (DAPI) staining was used adapting the protocol described by (Llop, Labella, et al. 2023). Samples (1 mL) were fixed with 1% formaldehyde for 20 minutes at room temperature, washed once with deionized water, and frozen at -20°C for 24 hours. Staining was performed using 0.25 mg/mL DAPI for 15 minutes at room temperature in the dark, followed by three washes with deionized water.

Micrographs were taken using a Zeiss LSM800 confocal laser scanning microscope. 5 μL of the stained samples were placed on 2% low-melting-point agarose pads. The microscopy settings were as follows: Ex 640 nm/em 650+ nm for auto-fluorescence, ex 405 nm/Em 470–617 nm for polyP, and ex 405 nm/em 470–617 nm for DAPI-control signals. Micrographs were coloured as red (auto-fluorescence) and light blue (polyP) to improve visualization contrast.

### Protein extraction, immunodetection, and band quantification

For protein extraction, 10 mL samples of *S. elongatus* liquid cultures at mid-exponential phase were collected at 0, 6 or 24 hours after addition of 1 mM IPTG. Cells were pelleted at 7300× g, flash frozen in liquid nitrogen, and stored at −20 °C. Pellets were resuspended in 60 μL of lysis buffer (10 mM Tris/HCl pH 7.5, 0.5 mM EDTA, 1mM β-mercaptoethanol, 1 mM phenylmethylsulfonyl fluoride (PMSF)), and cells were disrupted with 1 spoonful of 0.1 mm glass beads (≈30 µL) in a high-speed homogenizer Minibeadbeater (three pulses of 1 minute each, separated by 1-minute intervals on ice; speed of 5 m/s). Samples were centrifuged (5500× g for 5 min), and the supernatant fractions (crude protein extracts) were transferred to a new tube. Protein concentrations were estimated via the Bradford method (Bradford, 1976) using the PierceTM detergent-compatible Bradford assay kit (ThermoScientific, Waltham, MA, USA) in a VICTOR3TM 1420 Multilabel Plate Reader (PerkinElmer). Protein extracts were stored at −20 °C until needed.

For immunodetection, 50 μg of total protein extracts were loaded into a sodium dodecyl sulphate polyacrylamide gel (SDS-PAGE; 15% polyacrylamide). Electrophoresis was followed by immunoblotting onto 0.2 μm polyvinylidene fluoride membranes (from GE Healthcare Technologies, Inc., Chicago, IL, USA), and the membranes were subsequently blocked with 5% non-fat dry milk in phosphate-buffered saline with 0.1% Tween 20 (PBS-Tween) for 1 hour at room temperature. Membranes were then incubated overnight in PBS-Tween with 5% non-fat dried milk with a 1:300 dilution of PipY primary antibody (Labella et al. 2017) and for 1 h at room temperature with a 1:150,000 dilution of ECL rabbit IgG and an HRP-linked F(ab’)2 fragment (from a donkey, GE Healthcare). Visualization of bands was performed using SuperSignal WestFemto reagent (Thermo Fisher Scientific, Waltham, MA, USA) in a Biorad ChemiDoc Imager using the automatic exposure mode and avoiding pixel saturation.

### Computational methods

To measure cell length and protein band intensities images were analyzed using ImageJ v1.54g. The length was determined with the “Straight” tool of the program and the “Measure” button. The Western blot specific bands were selected using the “Rectangle” function, and their corresponding profiles were measured with the “Wand” tool.

To quantify the polyP signal, whole-cell fluorescence intensities from the polyP and DAPI-control channels in confocal images were measured using ImageJ v1.54g. PolyP intensities were normalized to the DAPI signal to account for differences in cell numbers and DAPI uptake.

To quantify *E. coli* colony sizes, Feret’s diameter was measured using ImageJ v1.54g. Images typed as “8-bit” were thresholded by Otsu’s method after background subtraction and subjected to a “watershed” process to separate colonies. The resultant images were adjusted to an ellipse and the particles filtered with a minimum size of 0.3 mm before measuring Feret’s diameter.

To perform statistical analyses, RStudio v2025.09.2 Build 418 was used (RStudio: Integrated Development for R. RStudio 2020).

To generate graphical representations of protein structures PyMOL v1.7.1.7 was used (The PyMOL Molecular Graphics System, Schrödinger, LLC).

## Results

### PipY mutations P63L or R210Q, but not K26A, prevent growth of *S. elongatus*

In contrast to PipY deficiency, PipY overexpression results in rather dramatic phenotypes (Labella et al. 2017; Llop, Labella, et al. 2023). Strain 1^S^Ptrc-PipY, used in the previous studies, carries an extra copy of *pipY* at the neutral site I (NSI) of the *S. elongatus* chromosome under the control of the IPTG-inducible promoter P*trc*. Tight control and inducible overexpression of PipY from the *pipY* gene is provided by the LacI repressor, also encoded within the NSI (see Fig. 2a).

**Figure 2.**
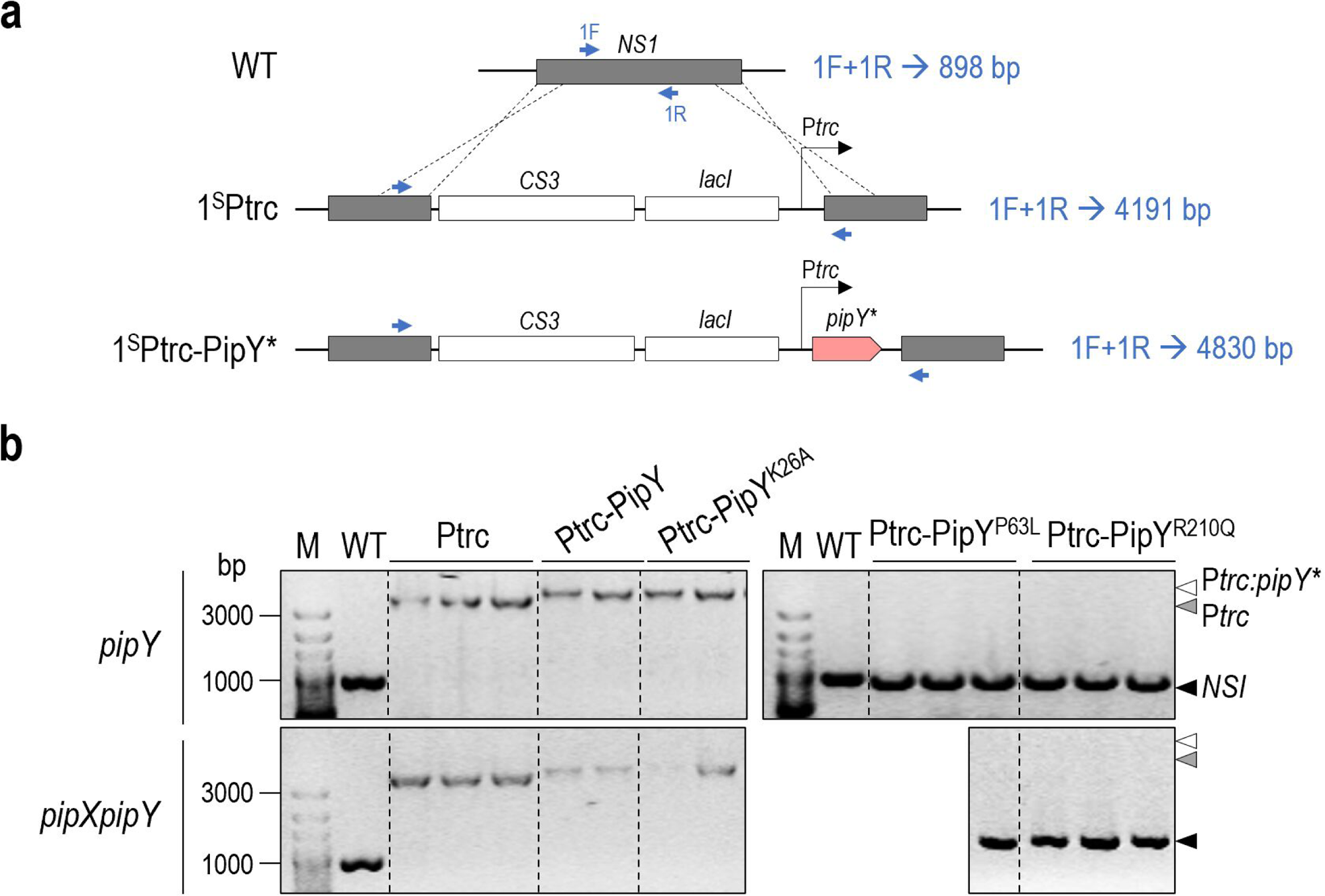
Constructs to overexpress PipY derivatives in *S. elongatus* and segregation analysis. **a)** Schematic representation of the *NSI* at the *S. elongatus* genomic region and of the constructs to express the PipY derivatives under the control of the *Ptrc* promoter in *pipY* or *pipXpipY* null backgrounds. Primer positions for PCR analysis are depicted by blue arrows, with the size of products on the right. **b)** Representative agarose gels showing PCR products obtained in the indicated strains and backgrounds after up to ten selective transfers, with the expected amplicon depicted by arrows on the right. M, 100 bp size marker. The number of colonies obtained in two independent transformations was >10^3^ for PipY or PipY^K26A^, and <15 for PipY^P63L^ or PipY^R210Q^, with no significative differences between both backgrounds. * refers to the WT, K26A, P63L, and R210Q versions of PipY.

To determine the effect of relevant point mutations on PipY activity in *S. elongatus* we first attempted construction of variants of strain 1^S^Ptrc-PipY, so the phenotypic effects of wild type and mutant PipY derivatives can be studied in parallel both at close to physiological levels and at very high levels of expression. With this in mind we first generated mutations encoding K26A, P63L or R210Q variants in plasmid pUAGC294, giving plasmids pUAGC1133, pUAGC1130 or pUAGC1129, respectively (Table 1). To avoid possible interference and/or gene conversion between the wild type and mutant alleles of *pipY,* we performed all experiments in *pipY* null strains. To explore a possible contribution of PipX to the phenotypes studied here while strengthening the robustness of the data we also obtained and studied *pipXpipY* derivatives in parallel by precisely replacing their ORF by the *cat* (chloramphenicol-acetyltransferase) gene as described in (Labella et al. 2017).

The *pipY* and *pipXpipY* mutants were next used as recipients in independent transformations with plasmids containing wild type, the three mutant alleles of *pipY* or, to provide a negative control for PipY activity, carrying no *pipY* gene (plasmid pUAGC280). Surprisingly, while constructs encoding PipY or PipY^K26A^ yield thousands of streptomycin-resistant transformants, the plasmids encoding PipY^P63L^ or PipY^R210Q^ produced just a few. In each case several transformants clones were PCR-analysed for the presence of the insert at NSI. For comparison the unmodified NSI was amplified from the WT strain.

As shown in Fig. 2b (left panel), PCR products of the expected size were found when the transformations involved 1^S^Ptrc-PipY, 1^S^Ptrc-PipY^K26A^ or 1^S^Ptrc constructs in both genetic backgrounds. In contrast, the PCR products of some of the few transformants obtained for 1^S^Ptrc-PipY^P63L^ and 1^S^Ptrc-PipY^R210Q^ constructs were indistinguishable from those of the WT control strain in the two genetic backgrounds (Fig. 2b, right panel). Since the unmodified *NSI* allele was the only one detected in those transformants, the results indicate that, even at relatively low levels, expression of PipY^P63L^ or PipY^R210Q^, but not of PipY^K26A^, prevents the appearance of *S. elongatus* colonies, independently of the *pipX* gene. Therefore, the results indicate that PipY^P63L^ or PipY^R210Q^, but not PipY^K26A^, behave as gain-of-function proteins that prevent growth of *S. elongatus* in our experimental conditions.

### Heterologous expression of PipY variants in *E. coli* parallels their behaviour in *S. elongatus*

The failure to obtain viable transformants expressing the PipY^P63L^ or PipY^R210Q^ variants (Fig. 2) raised questions on the mechanism(s) by which each of these proteins prevent *S. elongatus* growth. Since PipY is not required for *S. elongatus* viability, it follows that mutations P63L or R210Q are behaving as gain-of-function mutations and thus both PipY^P63L^ and PipY^R210Q^ must be interfering with cellular processes required for culture growth. To distinguish if this interference affects signalling pathways restricted to cyanobacteria or more universal processes conserved in phylogenetically distant bacterial groups, we next investigated the effect of expressing each of the PipY variants in *E. coli*.

It is worth noting that all plasmids constructed in this work were obtained in *E. coli* XL1-Blue and that this strain carries a *lacI^q^*gene to prevent expression of the cloned genes. Therefore, to unveil a possible phenotypic impact of the PipY variants in *E. coli* the LacI repressor must be inactivated. To this goal, plasmids pUAGC294, pUAGC1133, pUAGC1130 or pUAGC1129 were independently introduced into *E. coli* XL1-Blue and their corresponding transformants were selected in parallel in the absence or presence of the IPTG inducer.

As shown in Fig. 3a, multiple ampicillin-resistant transformants were obtained in all cases in the absence of IPTG. However, colonies derived from the plasmid bearing the P63L variant were noticeably smaller than those obtained with the other constructs (Fig. 3b), suggesting greater toxicity for PipY^P63L^ than for PipY^R210Q^. Since the latter protein is recovered at lower yield than the former from *E. coli* extracts (Tremiño et al. 2017), differences in expression level may, at least in part, explain their different toxicities. Upon IPTG addition (0.5 mM), the number of transformants remained unchanged only for the construct expressing PipY^K26A^ but decreased by 98% for PipY and by 100% for both PipY^P63L^ and PipY^R210Q^ (Fig. 3c). Independent corroboration of these results was next provided by transforming the same four plasmids into *E. coli* MG1655, a strain lacking the extra *lacI^q^*copy and therefore unable to provide additional repression of the promoter driving expression of the PipY variants (Fig. 3d). The substantially higher number of transformant obtained for PipY^K26A^ relative to PipY further supports that expression of PipY may also have a negative impact on *E. coli* viability that can be attenuated by the K26A mutation. No colonies were obtained when the proteins expressed were PipY^P63L^ or PipY^R210Q^, confirming the high toxicity of both variants in *E. coli*.

**Figure 3.**
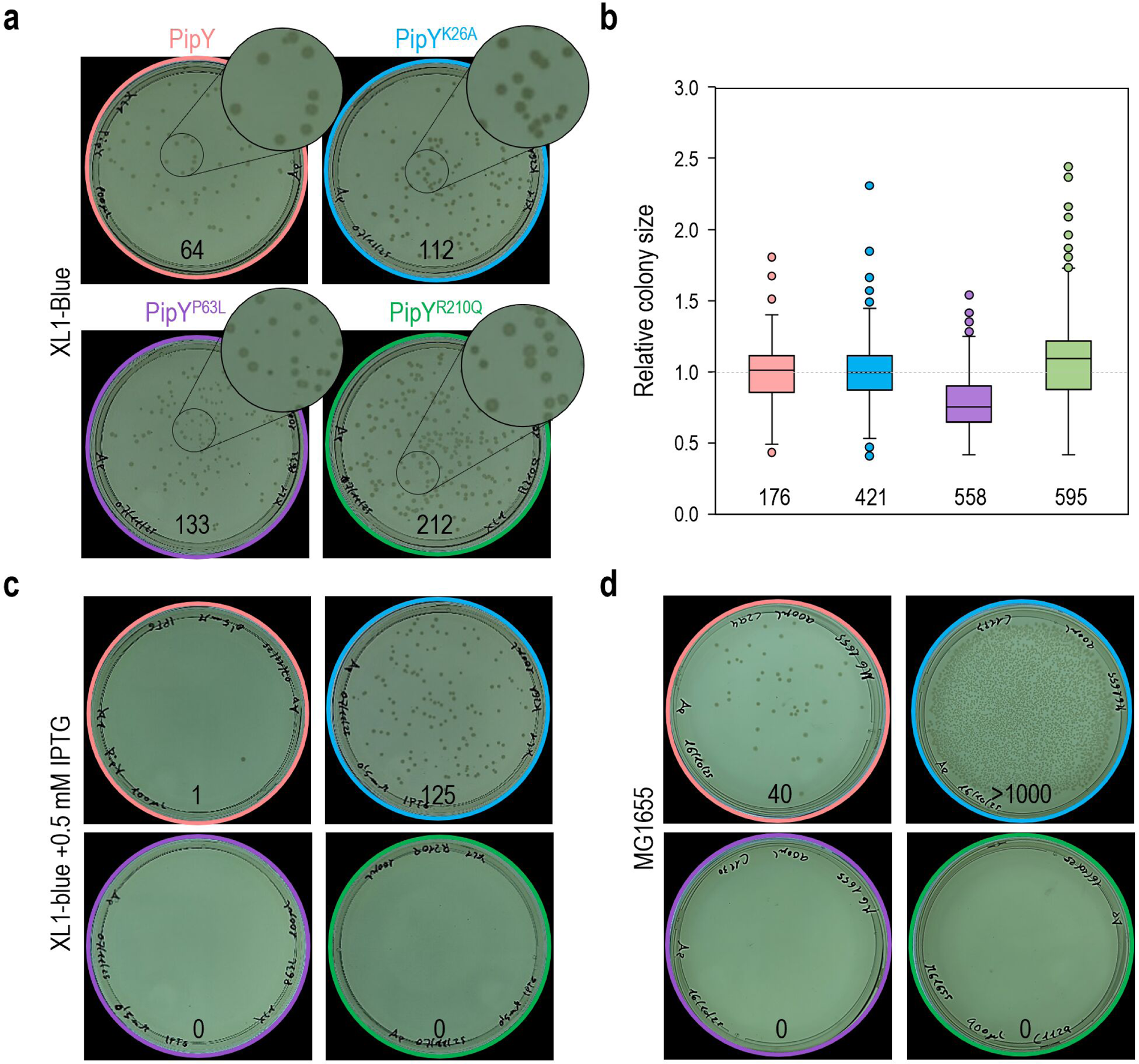
Effects of PipY variants in *E. coli* colony yield. Results of transformations with 50 ng of plasmids expressing the indicated PipY variants under the *Ptrc* promoter. **a)** XL1-blue strain (1/10 dilution). Insets show a 3× zoom of the indicated regions. **b)** Quantification of the Feret’s diameter of colonies, normalized to those of the PipY strain. The median, the interquartile range (box), and outliers (dots) are shown, along with the number of colonies measured. **c)** XL1-blue strain (1/10 dilution) in the presence of 0.5 mM IPTG. **d)** MG1655 strain (9/10 dilution). The number of colonies obtained is indicated inside each plate. Data is based on two independent experiments.

Taken together, our results show that the expression of PipY^R210Q^ and PipY^P63L^ variants inhibit growth in both *S. elongatus* and *E. coli*. Therefore, while the behaviour of PipY^K26A^ is consistent with loss-of-function (see also below), that of the “pathogenic” variants reveals gain-of-function in both bacterial model systems.

### Lys26 is required for PipY overexpression phenotypes on growth, chlorosis, cell size and overaccumulation of polyP

To gain insights into the factors affecting PipY overexpression phenotypes in *S. elongatus*, we addressed the impact of mutation K26A on previously studied traits, which include growth, chlorosis, cell size and overaccumulation of polyP (Llop, Labella, et al. 2023). The *S. elongatus* strains compared were *pipY* 1^S^Ptrc, *pipY* 1^S^Ptrc-PipY, *pipY* 1^S^Ptrc-PipY^K26A^, *pipXpipY* 1^S^Ptrc, *pipXpipY* 1^S^Ptrc-PipY, and *pipXpipY* 1^S^Ptrc-PipY^K26A^.

The impact on the different strains of IPTG induction on growth and pigment composition was followed by performing drop-plate assays or according to the visual appearance and absorbance spectra of liquid cultures (Fig. 4). As expected, no differences in growth were found amongst otherwise isogenic 1^S^Ptrc-PipY and 1^S^Ptrc strains in the absence of IPTG (left panel in Fig. 4a). However, addition of 50 μM IPTG prevented growth of 1^S^Ptrc-PipY but not of 1^S^Ptrc-PipY^K26A^ or 1^S^Ptrc strains (Fig. 4a, b), indicating that mutation K26A abrogates the inhibitory effects of PipY overexpression on growth.

**Figure 4.**
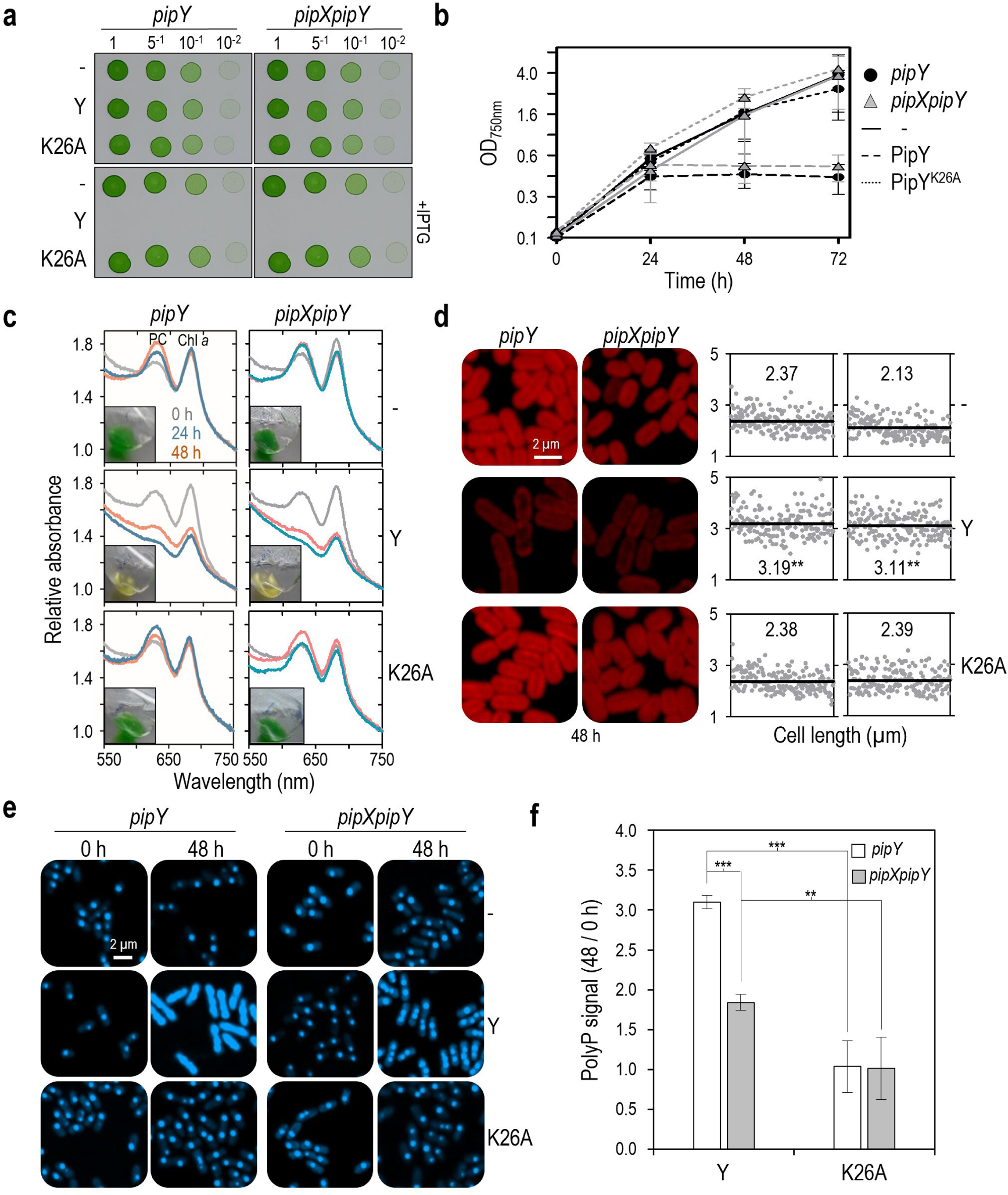
Effects of K26A mutation or *pipX* inactivation on PipY overexpression phenotypes in *S. elongatus*. Strains Ptrc (-) and derivatives expressing PipY (Y) or PipY^K26A^ (K26A) in the indicated backgrounds, grown in BG11 and, when indicated, supplemented with 50 µM IPTG. **a)** Drop-plate assay (5 μL; initial OD_750nm_= 0.5) with the dilutions indicated at the top. **b)** Growth curves (mean ± standard deviation). **c)** Whole absorbance spectra, indicating the peaks of phycocyanin (PC) and chlorophyll *a* (Chl *a*), and insets showing cultures after 48 h of IPTG induction. **d)** Representative confocal micrographs of autofluorescence signal (+40% brightness, –40% contrast; *left*) and scatterplot of cells length (n = 180 cells; *right*). Means are indicated by black lines and numbers inside the graphs. **e)** Representative confocal micrographs of DAPI-polyP channel. **f)** Ratio of the polyP signal (48 / 0 h) referred to the Ptrc strain (mean ± standard deviation). Pairwise comparisons using the Mann–Whitney *U* test with Bonferroni correction were performed. Significance levels were denoted as p ≤ 0.01 (**) or ≤0.001 (***). Effect size r was interpreted as large according to Cohen’s thresholds (r > 0.5). Data from two independent experiments are presented.

Next, we compared the dynamic of pigment loss induced by PipY overexpression in the different strains. As previously shown, addition of IPTG to *pipY* 1^S^Ptrc-PipY cultures triggered the yellow color and the progressive decrease in phycocyanin and chlorophyll *a* peaks characteristic of chlorosis (Fig. 4c, left panels). In contrast, *pipY* 1^S^Ptrc-PipY^K26A^ cultures maintained the normal green appearance and pigment composition throughout the 48 hours experiment, exactly as the negative control strain *pipY* 1^S^Ptrc.

The impact of IPTG induction on cell size was subsequently analysed by confocal microscopy at 0 and 48 h after IPTG induction (Fig. 4d, S1, S2). Representative micrographs showed that cell appearance and cell size were indistinguishable amongst the strains before IPTG addition (0), and that only 1^S^Ptrc-PipY cells were significantly longer in both backgrounds after IPTG addition (48 h; Fig. 4d). The lower autofluorescence of 1^S^Ptrc-PipY cells paralleled the loss of photosynthetic pigments shown in Fig. 4c. Scatter-plot representation of cell lengths for all six strains confirmed the significant increase in 1^S^Ptrc-PipY cells after IPTG induction (Fig. 4d, right).

The impact of IPTG induction on polyP accumulation, another relevant phenotype associated with PipY overexpression (Llop, Labella, et al. 2023), was subsequently analysed by confocal microscopy in the previous strains at 0 and 48 h after IPTG induction. Representative micrographs of DAPI-stained cells for the same six strains and two timepoints are shown in Fig. 4e. To quantify the polyP signal avoiding possible differences in cell number per field, the total signal in the DAPI–polyP channel (ex405, em470–617) was normalized against the corresponding DAPI channel (ex405, em410–470). No differences between IPTG treated or untreated cells were observed in strain 1^S^Ptrc-PipY^K26A^ (Fig. 4f), indicating that the K26A mutation abolished the over accumulation of polyP resulting from PipY overexpression.

In summary, all four PipY overexpression-related phenotypic features analyzed here are abolished by the K26A mutation, indicating that they require the PLP cofactor and/or the integrity of Lys26 at PipY.

### Inactivation of *pipX* affects overaccumulation of polyP but has no effect on other PipY overexpression phenotypes

Comparison between *pipY* and *pipXpipY* strains expressing PipY or PipY^K26A^ showed no phenotypic differences attributable to the presence or absence of *pipX* regarding growth, chlorosis or cell size (Fig. 4a,b,c,d). However, although *pipXpipY* 1^S^Ptrc-PipY cells showed significant accumulation of polyP afer 48h of induction, it was not as large as in *pipY* 1^S^Ptrc-PipY cells (Fig. 4e). PolyP signals were at least 1.7-fold lower from *pipXpipY* 1^S^Ptrc-PipY than from *pipY* 1^S^Ptrc-PipY cells (Fig. 4f), thus indicating that the *pipX* gene is required for the very high levels of polyP associated to PipY overexpression.

In summary, our results indicate that PipX plays a positive regulatory role in polyP accumulation, suggesting that both PipX and PipY may be required to increase polyP synthesis under certain environmental conditions. On the other hand, we cannot yet exclude regulation by PipX of the cellular targets involved in the other phenotypic traits analysed here in *S. elongatus* cells or cultures. Since we studied the effect of the presence or absence of *pipX* gene in conditions in which the PipX/PipY is abnormally low, a regulatory role of PipX in the studied traits could have been missed, particularly if it was a negative role.

### Mutation K26A increase the levels of PipY in *S. elongatus* in a PipX independent manner

Given that the phenotypes discussed above depend on PipY overaccumulation in *S. elongatus* ((Labella et al. 2017; Llop, Labella, et al. 2023); Fig. 4), it was important to exclude the possibility that mutation K26A impaired the stability of PipY therefore preventing accumulation of very high levels of PipY^K26A^ in the presence of IPTG. To investigate that possibility, samples of *pipY* 1^S^Ptrc-PipY, *pipXpipY* 1^S^Ptrc-PipY, *pipY* 1^S^Ptrc-PipY^K26A^, and *pipXpipY* 1^S^Ptrc-PipY^K26A^ strains were taken at 0, 6, and 24 h timepoints after IPTG induction, and subsequently analysed by Western blot with anti-PipY (Figs. 5 and S3).

**Figure 5.**
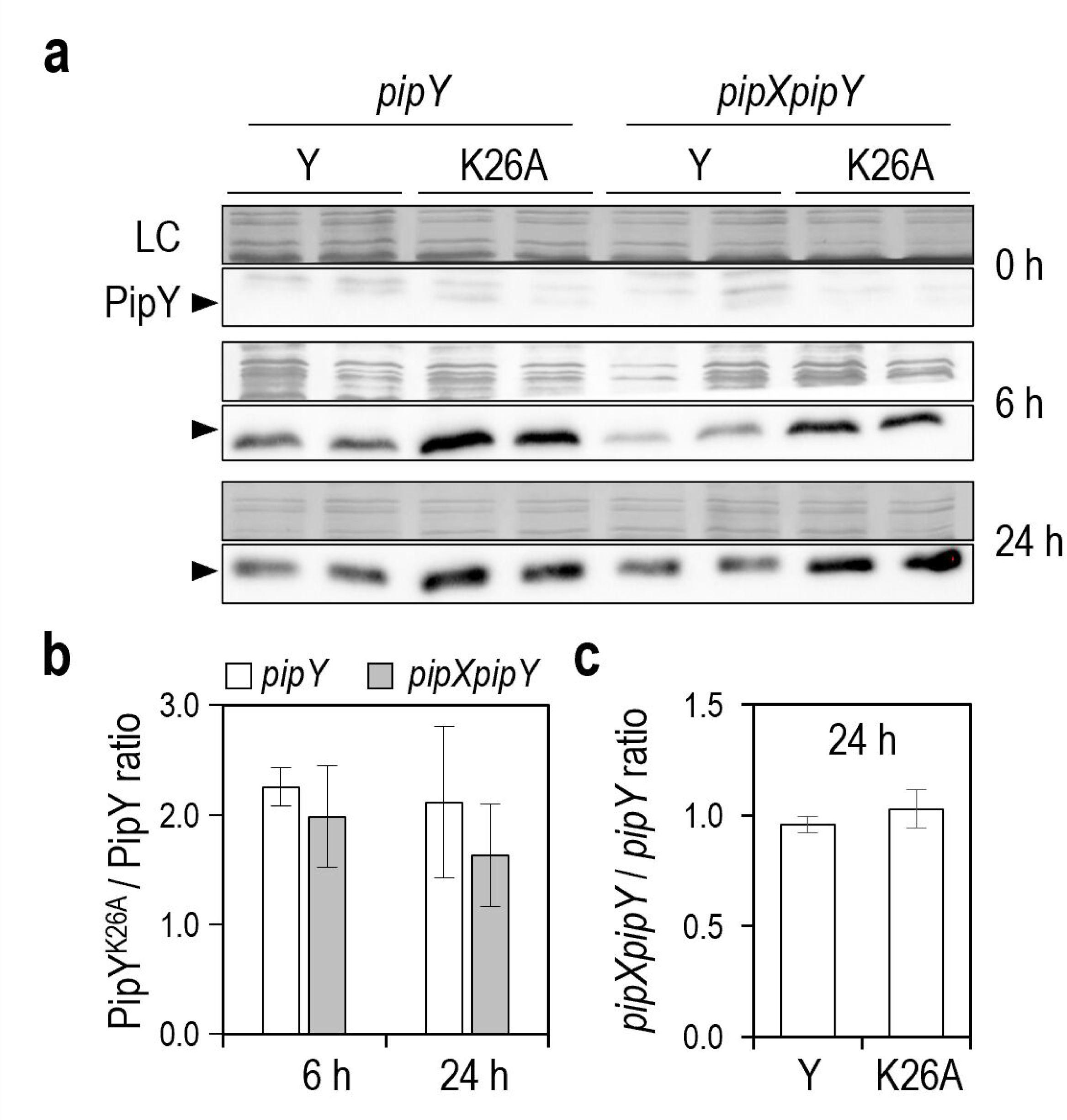
Effects of K26A mutation or *pipX* inactivation on PipY levels. **a)** Representative immunodetections (α-PipY) and loading control (LC) of Ptrc strains expressing PipY (Y) or PipY^K26A^ (K26A) in the indicated null mutant grew in the presence of 1 mM of IPTG. **b)** Ratio of PipY^K26A^/PipY protein levels, normalized to loading control. **c)** *pipXpipY/pipX* ratio at 24 h. Data are presented as means and error bars (standard deviation) from two biological replicates.

Although no differences between strains could be observed at timepoint 0, upon IPTG addition the signal from the PipY band was higher at both 6 and 24 h timepoints for PipY^K26A^ than for PipY (Fig. 5a). Quantification of band signals confirmed a ≈ 2-fold increase in PipY^K26A^ levels with respect to PipY (Fig. 5b) regardless of the presence or absence of the *pipX* gene (Fig. 5c).

Therefore, the failure of PipY^K26A^ to trigger any one of the PipY overexpression phenotypes in *S. elongatus* analysed here is not the result of lower accumulation of PipY^K26A^ in the presence of IPTG, indicating that either the PLP cofactor or Lys26 itself is required for PipY-triggered growth arrest, chlorosis, increased cell length and overaccumulation of polyP in *S. elongatus*.

## Discussion

In this work we have exploited the existing structural and functional information on the cyanobacterial model protein PipY to gain insights into the PLPBP family by focussing on the phenotypic effects conferred by PipY variants expressing point mutations that prevent PLP binding (K26A) or that mimic pathogenic changes (P63L and R210Q) found in the orthologous human protein.

The apparent correlation between the severity of *PLPBP* mutations causing vitamin B_6_-dependent epilepsy and the decreased ability of some of the corresponding PLPHP variants to bind the PLP cofactor (Johnstone et al. 2019) speaks of the importance of the PLP cofactor for PLPBP function in the homeostasis of B_6_ vitamers. For instances, both PipY^P63L^ and PLPHP^P87L^ were respectively less impaired than PipY^R210Q^ and PLPHP^R241Q^ *in vitro* (Tremiño et al. 2017; Tremiño et al. 2018) as well as in *E. coli*, where PLPHP^P87L^ but not PLPHP^R241Q^ complemented the pyridoxine-sensitivity of the *yggS* null mutant (Darin et al. 2016). It is worth noting that while the P63L mutation did not impair any of the parameters assayed at PipY, including protein yield, thermal stability, proper fold or PLP-content, the R210Q change resulted in a lower yield of the recombinant protein that, although well folded, contained hardly any PLP and showed decreased stability at high temperature. Pro63/87 is located at the tip of the turn between α3-β3, facing outwards and well away from the PLP cofactor, while Arg210/241 is an invariant residue that establish contacts with the PLP cofactor (Fig. 1). Considering that proteins with obvious defects in PLP binding were less functional *in vivo*, the prediction was that PipY^K26A^ would be at least as affected as PipY^R210Q^ when tested in *S. elongatus*.

Phenotypic analyses in *S. elongatus* indicated that mutation K26A caused complete loss of function. Despite the high levels of PipY^K26A^ protein achieved in *S. elongatus* (Fig. 5a, b) this protein did not confer any of the phenotypes associated with PipY overexpression, or indeed any other of the phenotypic features analysed in *S. elongatus*. Furthermore, while heterologous expression of PipY decreased viability in *E. coli*, expression of PipY^K26A^ did not, indicating that the differential behaviour of wild type and mutant proteins is maintained across bacterial species. Since the equivalent mutation (K36A) at the *E. coli* homolog YggS also increases the stability of the protein (Tramonti et al. 2022), our results emphasise the importance and common roles of this invariant lysine within bacterial PLPBPs. Not surprisingly, PipX, a protein unique to cyanobacteria, appears to play no role in the different stability of PipY and PipY^K26A^ proteins (Fig. 5c).

Interestingly, the PipY^R210Q^ variant, which is also impaired in PLP binding (Tremiño et al. 2017), did not decrease PipY toxicity. On the contrary, it resulted toxic in both *S. elongatus* and *E. coli* (Figs. 2 and 3) and thus the opposite phenotypes caused by two mutations abolishing or drastically impairing PLP binding argues against PipY toxicity requiring the presence of the PLP cofactor. It is thus likely that toxicity depends on a particular protein conformation which could not be adopted by PipY^K26A^. Since our results indicate that Lys26 must have additional functions beyond cofactor binding, it is reasonable to think that PipY must interact with yet unknown targets in bacterial cells. In this context, the simplest interpretation for the opposite phenotypes of PipY^K26A^ and PipY^R210Q^ is that Lys26, and not PLP itself, was required for PipY toxicity and that mutation K26A would likely impair interactions with those hypothetical targets while R210Q, as well as P63L, would favour it.

Since two mutations (P63L and R210Q) that have different effects on the affinity for PLP confer the same gain-of-function phenotype, increasing PipY toxicity in both *S. elongatus* and *E. coli* cells even when expressed at very low levels (Figs. 2 and 3), the PLP cofactor would not be directly involved in PipY toxicity. Instead, the results suggested that PipY interacts with yet unknown targets in both bacterial systems and that the PipY^P63L^ and PipY^R210Q^ variants do it to greater extent.

Since PipY^P63L^ conferred higher toxicity than PipY^R210Q^ in bacteria (Fig. 3), it appears that the P63L mutation confers less severe loss-of-function and more severe gain-of-function phenotypes than R210Q for respectively, PLP-dependent and PLP-independent activities. These results, in complete agreement with the higher protein yield of PipY^P63L^ found in *E. coli* (Tremiño et al. 2017), support the idea that for PLP-independent functions, that is, toxicity, there would be two alternative conformations of relevance, one represented by PipY^K26A^ (inactive) and the other one represented by PipY^P63L^ or PipY^R210Q^ (active). Taken together, the results suggest that in addition to vitamer B_6_ homeostasis PLPBP is involved in PLP-independent activities.

Concerning the nature of PipY interactants, a recent work shows that PLPBP members are RNA binding proteins and that the apo-forms of YggS and human PLPHP bind RNA with much greater affinity than the holo-forms or the YggS^K36A^ variant, which adopts a holo-like conformation despite the absence of PLP (Graziani et al. 2024). According to that work, PLP would have a role as an effector molecule inhibiting the binding of RNA to PLPBPs (Graziani et al. 2024). It is that tempting to propose that PipY toxicity is related to its ability to bind RNA and that the apo-form is involved.

A comparative analysis of the PipY structures from *S. elongatus* (PLP-bound PipY, PDB: 5NM8, versus apo PipY, PDB: 5NLC) reveal changes associated with PLP binding at helices α1, α2, α6, and α9, as well as the β5–α6, β6–α7, and β7–α8 loops (Fig. 6a,b,c; (Tremiño et al. 2017)). Use of the KVFinder web tool (https://kvfinder-web.cnpem.br/; (Guerra et al. 2020; Guerra et al. 2023)) detected differences in the solvent-accessible PLP-binding cavity, with its volume increasing from 144.94 Å³ to 161.57 Å³ upon PLP release. Thus, close and open conformations of the protein surface can be respectively associated to the holo or apo forms.

**Figure 6.**
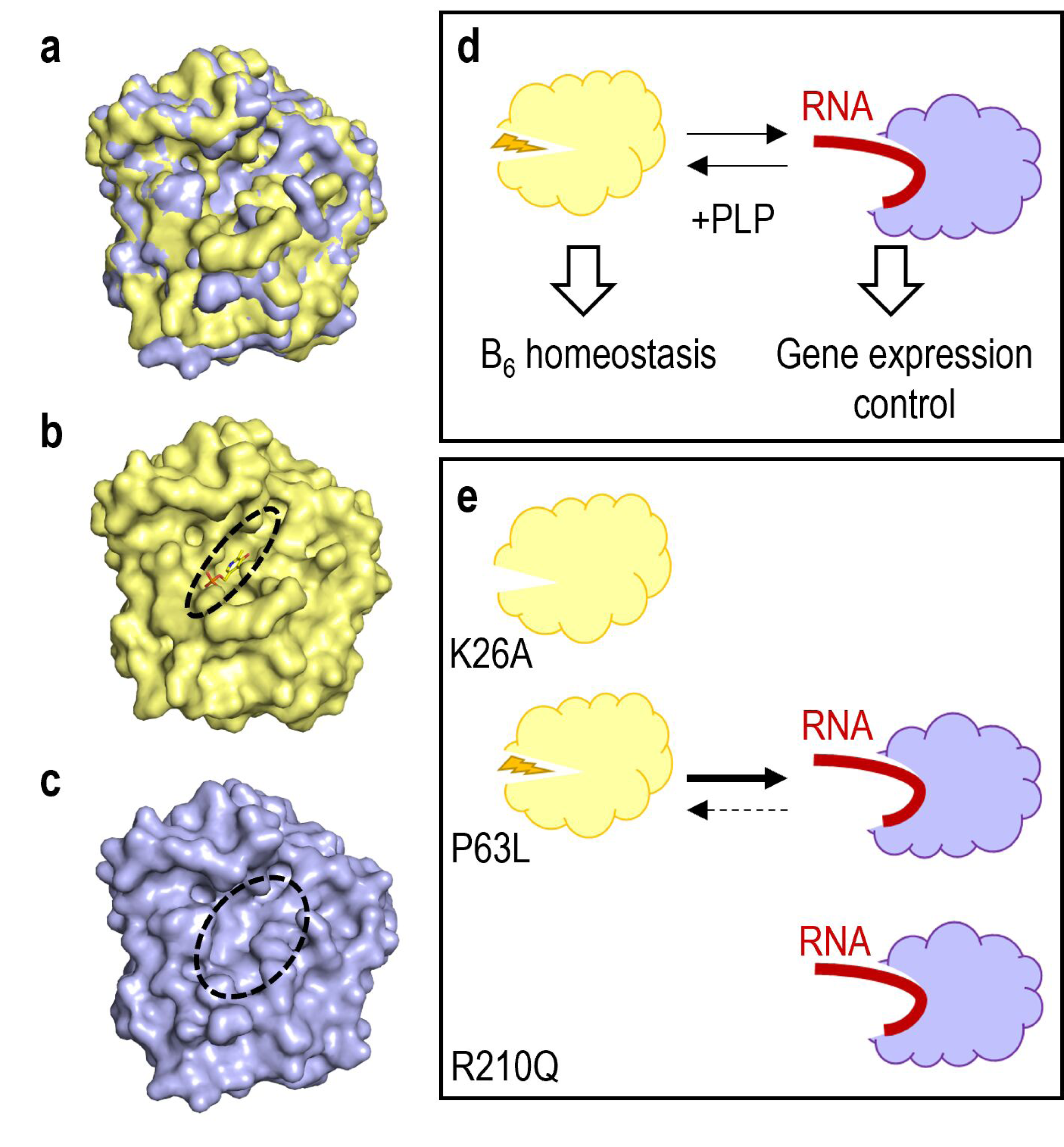
PLP driven changes and model for PipY functions based on alternative conformations. **a)** Superposition of PipY-apo (blue) onto PipY-PLP (yellow). **b)** PipY–PLP. **c)** PipY-apo. The dashed ellipses mark solvent-accessible regions. **d)** Schematic representation of the holo and apo forms bound to their respective ligands, with indication of functions. **e)** Predicted effect of the indicated mutations on protein conformation. See text for additional details.

A working model for PipY functions is schematically illustrated in Fig. 6d,e. We propose that like YggS^K36A^, PipY^K26A^ would adopt a holo-like conformation with low affinity for RNA and that in the case of PipY^R210Q^, insufficient binding of the PLP cofactor would displace the equilibrium towards the apo-form. Since PipY^P63L^ is not impaired in PLP binding, displacement of the equilibrium towards the apo-form of PipY by mutation P63L requires a different explanation. We speculate that Pro63 may play a role in maintaining the holo-form by imposing rigidity to the α3-β3 loop. Based on the different expression levels for PipY^R210Q^ and PipY^P63L^ (Tremiño et al. 2017), the prediction is that *in vivo* virtually all of PipY^R210Q^ would be in the apo-form, while for the more abundant protein PipY^P63L^ there would still be enough amount of the minoritarian holo form to explain its function on vitamer B_6_ homeostasis. In support of the model, the program GPSite (https://bio-web1.nscc-gz.cn/app/GPSite; (Yuan et al. 2024)) predicts that mutation K26A, but not R210Q or P63L, impair RNA binding (Table S2).

Last but not least, transcriptomic and proteomic changes associated to PLPBP deficient cells (Labella et al. 2017; Fux and Sieber 2020) support the recent proposal that PLPBP may be part of a regulatory mechanism of translation linked to amino acid levels (Graziani et al. 2024). Since PipX, a multifunctional protein co-expressed with PipY, is also involved in carbon/nitrogen homeostasis, transcription and translation regulation (Llácer et al. 2010; Espinosa et al. 2014; Labella et al. 2016; Labella et al. 2017; Cantos et al. 2019; Jerez et al. 2021), understanding the functions of the *pipXY* operon in cyanobacteria should help address the universal functions of PLPBP.

## Data availability statement

The original contributions presented in the study are included in the article/Supplementary material. Further inquiries can be directed to the corresponding author.

## Acknowledgments

The authors thank S. Bibak for Western blots, R. Cantos, T. Mata-Balaguer, P. Salinas, and L. Fuertes-García for technical contributions or advice.

## Conflict of interest

The authors declare that the research was conducted in the absence of any commercial or financial relationships that could be construed as a potential conflict of interest.

Alternative conformations of pyridoxal-phosphate binding proteins explain phenotypic effects of point mutations in cyanobacteria and humans

---

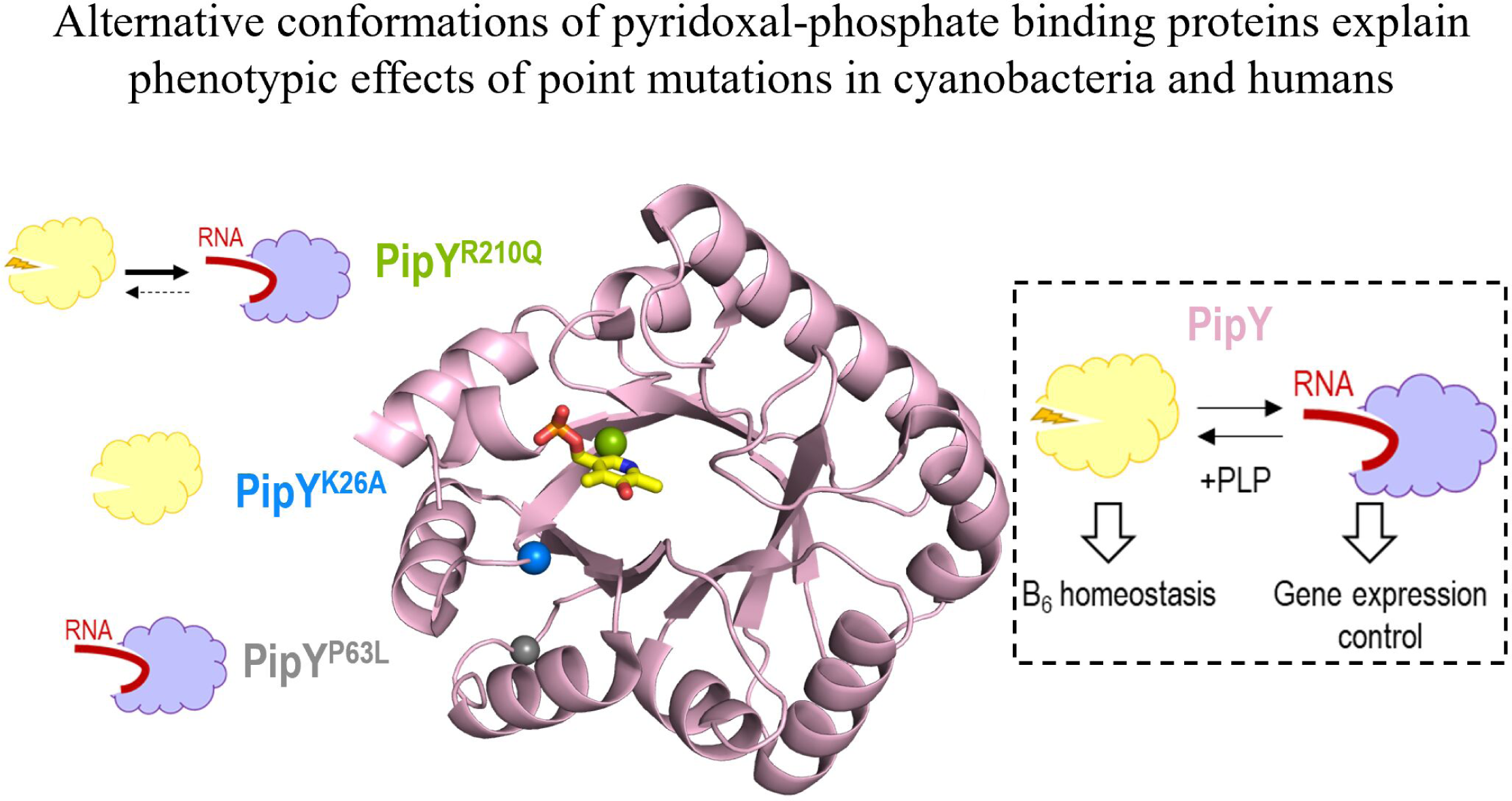

## References

Bullock WO, Fernandez JM, Short JM. 1987. XL1-Blue—a high-efficiency plasmid transforming *recA Escherichia coli* strain with β-galactosidase selection. Biotechniques. 5(3):376–379.

Burillo S, Luque I, Fuentes I, Contreras A. 2004. Interactions between the Nitrogen Signal Transduction Protein PII and *N* -Acetyl Glutamate Kinase in Organisms That Perform Oxygenic Photosynthesis. J Bacteriol. 186(11):3346–3354. doi:10.1128/JB.186.11.3346-3354.2004.

Cantos R, Labella JI, Espinosa J, Contreras A. 2019. The nitrogen regulator PipX acts *in cis* to prevent operon polarity. Environ Microbiol Rep. 11(4):495–507. doi:10.1111/1758-2229.12688.

Darin N, Reid E, Prunetti L, Samuelsson L, Husain RA, Wilson M, El Yacoubi B, Footitt E, Chong WK, Wilson LC, et al. 2016. Mutations in *PROSC* Disrupt Cellular Pyridoxal Phosphate Homeostasis and Cause Vitamin-B_6_-Dependent Epilepsy. The American Journal of Human Genetics. 99(6):1325–1337. doi:10.1016/j.ajhg.2016.10.011.

Espinosa J, Castells MA, Laichoubi KB, Contreras A. 2009. Mutations at *pipX* Suppress Lethality of P II -Deficient Mutants of *Synechococcus elongatus* PCC 7942. J Bacteriol. 191(15):4863–4869. doi:10.1128/JB.00557-09.

Espinosa J, Forchhammer K, Burillo S, Contreras A. 2006. Interaction network in cyanobacterial nitrogen regulation: PipX, a protein that interacts in a 2-oxoglutarate dependent manner with PII and NtcA. Mol Microbiol. 61(2):457–469. doi:10.1111/j.1365-2958.2006.05231.x.

Espinosa J, Forchhammer K, Contreras A. 2007. Role of the *Synechococcus* PCC 7942 nitrogen regulator protein PipX in NtcA-controlled processes. Microbiology (N Y). 153(3):711–718. doi:10.1099/mic.0.2006/003574-0.

Espinosa J, Labella JI, Cantos R, Contreras A. 2018. Energy drives the dynamic localization of cyanobacterial nitrogen regulators during diurnal cycles. Environ Microbiol. 20(3):1240–1252. doi:10.1111/1462-2920.14071.

Espinosa J, Rodríguez-Mateos F, Salinas P, Lanza VF, Dixon R, de la Cruz F, Contreras A. 2014. PipX, the coactivator of NtcA, is a global regulator in cyanobacteria. Proceedings of the National Academy of Sciences. 111(23):201404030–201404097. doi:10.1073/pnas.1404097111.

Eswaramoorthy S, Gerchman S, Graziano V, Kycia H, Studier FW, Swaminathan S. 2003. Structure of a yeast hypothetical protein selected by a structural genomics approach. Acta Crystallogr D Biol Crystallogr. 59(1):127–135. doi:10.1107/S0907444902018012.

Farkas P, Fitzpatrick TB. 2024. Two pyridoxal phosphate homeostasis proteins are essential for management of the coenzyme pyridoxal 5′-phosphate in *Arabidopsis*. Plant Cell. 36(9):3689–3708. doi:10.1093/plcell/koae176.

Forcada-Nadal A, Bibak S, Salinas P, Contreras A, Rubio V, Llácer JL. 2025. Structures of the cyanobacterial nitrogen regulators NtcA and PipX complexed to DNA shed light on DNA binding by NtcA and implicate PipX in the recruitment of RNA polymerase. Nucleic Acids Res. 53(4). doi:10.1093/nar/gkaf096.

Forchhammer K, Schwarz R. 2019. Nitrogen chlorosis in unicellular cyanobacteria – a developmental program for surviving nitrogen deprivation. Environ Microbiol. 21(4):1173–1184. doi:10.1111/1462-2920.14447.

Froger A, Hall JE. 2007. Transformation of Plasmid DNA into *E. coli* Using the Heat Shock Method. Journal of Visualized Experiments.(6). doi:10.3791/253.

Fux A, Sieber SA. 2020. Biochemical and Proteomic Studies of Human Pyridoxal 5′-Phosphate-Binding Protein (PLPBP). ACS Chem Biol. 15(1):254–261. doi:10.1021/acschembio.9b00857.

Giner-Lamia J, Robles-Rengel R, Hernández-Prieto MA, Muro-Pastor MI, Florencio FJ, Futschik ME. 2017. Identification of the direct regulon of NtcA during early acclimation to nitrogen starvation in the cyanobacterium *Synechocystis* sp. PCC 6803. Nucleic Acids Res. 45(20):11800–11820. doi:10.1093/nar/gkx860.

Graziani C, Barile A, Parroni A, di Salvo ML, De Cecio I, Colombo T, Babor J, de Crécy-Lagard V, Contestabile R, Tramonti A. 2024. The ubiquitous pyridoxal 5′-phosphate-binding protein is also an RNA-binding protein. Protein Science. 33(12). doi:10.1002/pro.5242.

Guerra JV da S, Ribeiro Filho HV, Bortot LO, Honorato RV, Pereira JG de C, Lopes-de-Oliveira PS. 2020. ParKVFinder: A thread-level parallel approach in biomolecular cavity detection. SoftwareX. 12:100606. doi:10.1016/j.softx.2020.100606.

Guerra JVS, Ribeiro-Filho H V, Pereira JGC, Lopes-de-Oliveira PS. 2023. KVFinder-web: a web-based application for detecting and characterizing biomolecular cavities. Nucleic Acids Res. 51(W1):W289–W297. doi:10.1093/nar/gkad324.

Guyer MS, Reed RR, Steitz JA, Low KB. 1981. Identification of a Sex-factor-affinity Site in *E. coli* as gamma delta. Cold Spring Harb Symp Quant Biol. 45(0):135–140. doi:10.1101/SQB.1981.045.01.022.

He S, Chen Y, Wang L, Bai X, Bu T, Zhang J, Lu M, Ha N-C, Quan C, Nam KH, et al. 2022. Structural and Functional Analysis of the Pyridoxal Phosphate Homeostasis Protein YggS from *Fusobacterium nucleatum*. Molecules. 27(15):4781. doi:10.3390/molecules27154781.

Ito T. 2022. Role of the conserved pyridoxal 5ʹ-phosphate-binding protein YggS/PLPBP in vitamin B_6_ and amino acid homeostasis. Biosci Biotechnol Biochem. 86(9):1183–1191. doi:10.1093/bbb/zbac113.

Ito T, Hori R, Hemmi H, Downs DM, Yoshimura T. 2020. Inhibition of glycine cleavage system by pyridoxine 5′-phosphate causes synthetic lethality in *glyA yggS* and *serA yggS* in *Escherichia coli*. Mol Microbiol. 113(1):270–284. doi:10.1111/mmi.14415.

Ito T, Yamamoto K, Hori R, Yamauchi A, Downs DM, Hemmi H, Yoshimura T. 2019. Conserved Pyridoxal 5’-Phosphate-Binding Protein YggS Impacts Amino Acid Metabolism through Pyridoxine 5’-Phosphate in *Escherichia coli*. Zhou N-Y, editor. Appl Environ Microbiol. 85(11):NA-NA. doi:10.1128/AEM.00430-19.

Jerez C, Llop A, Salinas P, Bibak S, Forchhammer K, Contreras A. 2024. Analysing the Cyanobacterial PipX Interaction Network Using NanoBiT Complementation in *Synechococcus elongatus* PCC7942. Int J Mol Sci. 25(9):4702. doi:10.3390/ijms25094702.

Jerez C, Salinas P, Llop A, Cantos R, Espinosa J, Labella JI, Contreras A. 2021. Regulatory Connections Between the Cyanobacterial Factor PipX and the Ribosome Assembly GTPase EngA. Front Microbiol. 12(NA):781760-NA. doi:10.3389/fmicb.2021.781760.

Johnstone DL, Al-Shekaili HH, Tarailo-Graovac M, Wolf NI, Ivy AS, Demarest S, Roussel Y, Ciapaite J, van Roermund CWT, Kernohan KD, et al. 2019. PLPHP deficiency: clinical, genetic, biochemical, and mechanistic insights. Brain. 142(3):542–559. doi:10.1093/brain/awy346.

Labella JI, Cantos R, Espinosa J, Forcada-Nadal A, Rubio V, Contreras A. 2017. PipY, a Member of the Conserved COG0325 Family of PLP-Binding Proteins, Expands the Cyanobacterial Nitrogen Regulatory Network. Front Microbiol. 8(NA):1244. doi:10.3389/fmicb.2017.01244.

Labella JI, Cantos R, Salinas P, Espinosa J, Contreras A. 2020. Distinctive Features of PipX, a Unique Signaling Protein of Cyanobacteria. Life. 10(6):79. doi:10.3390/life10060079.

Labella JI, Obrebska A, Espinosa J, Salinas P, Forcada-Nadal A, Tremiño L, Rubio V, Contreras A. 2016. Expanding the Cyanobacterial Nitrogen Regulatory Network: The GntR-Like Regulator PlmA Interacts with the PII-PipX Complex. Front Microbiol. 7(NA):1677. doi:10.3389/fmicb.2016.01677.

Laichoubi KB, Beez S, Espinosa J, Forchhammer K, Contreras A. 2011. The nitrogen interaction network in *Synechococcus* WH5701, a cyanobacterium with two PipX and two PII-like proteins. Microbiology (NY). 157(4):1220–1228. doi:10.1099/mic.0.047266-0.

Laichoubi KB, Espinosa J, Castells MA, Contreras A. 2012. Mutational Analysis of the Cyanobacterial Nitrogen Regulator PipX. Janssen PJ, editor. PLoS One. 7(4):e35845. doi:10.1371/journal.pone.0035845.

Llácer JL, Espinosa J, Castells MA, Contreras A, Forchhammer K, Rubio V. 2010. Structural basis for the regulation of NtcA-dependent transcription by proteins PipX and PII. Proceedings of the National Academy of Sciences. 107(35):15397–15402. doi:10.1073/pnas.1007015107.

Llop A, Bibak S, Cantos R, Salinas P, Contreras A. 2023. The ribosome assembly GTPase EngA is involved in redox signaling in cyanobacteria. Front Microbiol. 14(NA):1242616-NA. doi:10.3389/fmicb.2023.1242616.

Llop A, Labella JI, Borisova M, Forchhammer K, Selim KA, Contreras A. 2023. Pleiotropic effects of PipX, PipY, or RelQ overexpression on growth, cell size, photosynthesis, and polyphosphate accumulation in the cyanobacterium *Synechococcus elongatus* PCC7942. Front Microbiol. 14(NA):1141775-NA. doi:10.3389/fmicb.2023.1141775.

Moronta-Barrios F, Espinosa J, Contreras A. 2013. Negative control of cell size in the cyanobacterium *Synechococcus elongatus* PCC 7942 by the essential response regulator RpaB. FEBS Lett. 587(5):504–509. doi:10.1016/j.febslet.2013.01.023.

Prunetti L, El Yacoubi B, Schiavon CR, Kirkpatrick E, Huang L, Bailly M, El Badawi-Sidhu M, Harrison K, Gregory JF, Fiehn O, et al. 2016. Evidence that COG0325 proteins are involved in PLP homeostasis. Microbiology (N Y). 162(4):694–706. doi:10.1099/mic.0.000255.

Rippka R, Deruelles J, Waterbury JB, Herdman M, Stanier RY. 1979. Generic Assignments, Strain Histories and Properties of Pure Cultures of Cyanobacteria. Microbiology (N Y). 111(1):1–61. doi:10.1099/00221287-111-1-1.

RStudio: Integrated Development for R. RStudio. 2020. [accessed 2025 Jan 1]. http://www.rstudio.com/.

Salinas P, Bibak S, Cantos R, Tremiño L, Jerez C, Mata T, Contreras A. 2024. Studies on the PII-PipX-NtcA Regulatory Axis of Cyanobacteria Provide Novel Insights into the Advantages and Limitations of Two-Hybrid Systems for Protein Interactions. Int J Mol Sci. doi: 10.3390/ijms25105429

Schneider G, Käck H, Lindqvist Y. 2000. The manifold of vitamin B_6_ dependent enzymes. Structure. 8(1):R1–R6. doi:10.1016/S0969-2126(00)00085-X.

Schwarz R, Grossman AR. 1998. A response regulator of cyanobacteria integrates diverse environmental signals and is critical for survival under extreme conditions. Proceedings of the National Academy of Sciences. 95(18):11008–11013. doi:10.1073/pnas.95.18.11008.

Spät P, Klotz A, Rexroth S, Maček B, Forchhammer K. 2018. Chlorosis as a Developmental Program in Cyanobacteria: The Proteomic Fundament for Survival and Awakening. Molecular & Cellular Proteomics. 17(9):1650–1669. doi:10.1074/mcp.RA118.000699.

Taton A, Erikson C, Yang Y, Rubin BE, Rifkin SA, Golden JW, Golden SS. 2020. The circadian clock and darkness control natural competence in cyanobacteria. Nat Commun. 11(1):1688. doi:10.1038/s41467-020-15384-9.

Tramonti A, Ghatge MS, Babor JT, Musayev FN, di Salvo ML, Barile A, Colotti G, Giorgi A, Paredes SD, Donkor AK, et al. 2022. Characterization of the *Escherichia coli* pyridoxal 5′-phosphate homeostasis protein (YggS): Role of lysine residues in PLP binding and protein stability. Protein Science. 31(11):e4471-NA. doi:10.1002/pro.4471.

Tremiño L, Forcada-Nadal A, Contreras A, Rubio V. 2017. Studies on cyanobacterial protein PipY shed light on structure, potential functions, and vitamin B_6_ -dependent epilepsy. FEBS Lett. 591(20):3431–3442. doi:10.1002/1873-3468.12841.

Tremiño L, Forcada-Nadal A, Rubio V. 2018. Insight into vitamin B_6_ -dependent epilepsy due to *PLPBP* (previously *PROSC*) missense mutations. Hum Mutat. 39(7):1002–1013. doi:10.1002/humu.23540.

Tremiño L, Llop A, Rubio V, Contreras A. 2022. The Conserved Family of the Pyridoxal Phosphate-Binding Protein (PLPBP) and Its Cyanobacterial Paradigm PipY. Life. 12(10):1622. doi:10.3390/life12101622.

Vu HN, Ito T, Downs DM. 2020. The Role of YggS in Vitamin B_6_ Homeostasis in *Salmonella enterica* Is Informed by Heterologous Expression of Yeast *SNZ3*. Metcalf WW, editor. J Bacteriol. 202(22):NA-NA. doi:10.1128/JB.00383-20.

Yuan Q, Tian C, Yang Y. 2024. Genome-scale annotation of protein binding sites via language model and geometric deep learning. Elife. 13. doi:10.7554/eLife.93695.

